# CREB-mediated transcriptional activation of NRMT1 drives muscle differentiation

**DOI:** 10.1101/2021.05.24.445473

**Authors:** John G. Tooley, James P. Catlin, Christine E. Schaner Tooley

## Abstract

The N-terminal methyltransferase NRMT1 is an important regulator of protein-DNA interactions and plays a role in many cellular processes, including mitosis, cell cycle progression, chromatin organization, DNA damage repair, and transcriptional regulation. Accordingly, loss of NRMT1 results in both developmental pathologies and oncogenic phenotypes. Though NRMT1 plays such important and diverse roles in the cell, little is known about its own regulation. To better understand the mechanisms governing NRMT1 expression, we first identified its predominant transcriptional start site and minimal promoter region with predicted transcription factor motifs. We then used a combination of luciferase and binding assays to confirm CREB1 as the major regulator of NRMT1 transcription. We tested which conditions known to activate CREB1 also activated NRMT1 transcription, and found CREB1-mediated NRMT1 expression was increased during recovery from serum starvation and muscle cell differentiation. To determine how NRMT1 expression affects myoblast differentiation, we used CRISPR/Cas9 technology to knock out NRMT1 expression in immortalized C2C12 mouse myoblasts. C2C12 cells depleted of NRMT1 lacked Pax7 expression and were unable to proceed down the muscle differentiation pathway. Instead, they took on characteristics of C2C12 cells that have transdifferentiated into osteoblasts, including increased alkaline phosphatase and type I collagen expression and decreased proliferation. These data implicate NRMT1 as an important downstream target of CREB1 during muscle cell differentiation.

## Introduction

The N-terminal methyltransferase NRMT1 (N-terminal RCC1 methyltransferase 1) was discovered just over a decade ago, but in this short time has been shown to play an important role in many biological processes ^1–3^. NRMT1 has more than 300 predicted targets ^4^ and many significant verified targets including regulator of chromosome condensation 1 (RCC1), retinoblastoma protein (RB), the oncoprotein SET, DNA damage-binding protein 2 (DDB2), and centromere proteins A and B (CENP-A and CENP-B) ^1,5–7^. N-terminal methylation regulates protein/DNA interactions and loss of NRMT1 results in reduced interaction of its substrates with DNA ^5,6,8^.

The phenotypic consequences of NRMT1 loss include multi-polar mitotic spindles, impaired DNA damage repair, altered chromatin structure, and transcriptional misregulation ^1,5,9^. NRMT1 has been shown to act as a tumor suppressor in breast cancer cells, as its loss promotes cell proliferation, cell migration, colony formation, and xenograft growth ^2^. NRMT1 also plays important developmental roles, as only 30% of NRMT1 knockout mice (*Nrmt1^-/-^*) are viable past 6 months of age, and those that live exhibit premature ageing phenotypes and neurodegeneration ^3,10^. We have recently shown that their age-dependent neurodegeneration is preceded by mis-regulation of the neural stem cell (NSC) population ^10^. The postnatal quiescent NSC pool initiates premature proliferation and differentiation but the resulting neurons are unable to leave the cell cycle and initiate terminal differentiation, ultimately leading to apoptosis ^10^.

Though NRMT1 has been identified as an important developmental and oncogenic regulator, very little is known about its own regulation. Global proteomic studies have identified a handful of NRMT1 post-translational modifications (PTMs), including phosphorylation, arginine methylation, ubiquitination, and sumoylation ^11–15^. While it is not yet clear how any of these protein modifications affect NRMT1 function, it has recently been shown that the translational efficiency of NRMT1 is regulated by m6A RNA methylation ^16^. The transcriptional machinery regulating NRMT1 expression also remains elusive. NRMT1 has near ubiquitous expression in both human and mouse tissue ^17–19^, but its mRNA levels fluctuate during embryogenesis and development, indicating it does have dynamic transcriptional regulation ^20,21^. Here we identify cAMP-response element binding protein 1 (CREB1) as the first known transcriptional regulator of NRMT1.

CREB1 is an important regulator of many biological processes. It is activated through phosphorylation at Ser 133 by multiple cellular signaling cascades, including Ras/Raf/MAPK/p90RSK, Ca^2+/^CaMK, PI3K/Akt/GSK3β, and cAMP/PKA ^22^. The phosphorylated form of CREB1 (pCREB1) is able to bind cAMP response elements (CRE) in the promoter regions of its target genes, recruit CREB-binding protein (CBP), and initiate transcription ^23^. CREB1 phosphorylation is an important regulator of DNA damage repair and is activated in response to oxidative stress ^24^ and ionizing radiation ^25^. It is also a very important regulator of cell cycle and is activated during differentiation of many stem cell populations, including hematopoietic ^26^, mesenchymal ^27^, and neural stem cells ^28^.

Here we establish that CREB1-mediated transcription of NRMT1 is activated during both recovery from serum starvation and myoblast differentiation, and this activation has functional consequences. C2C12 mouse myoblasts depleted of NRMT1 fail to express Paired Box 7 (Pax7), a master regulator of postnatal muscle growth and regeneration ^29^, or any other downstream regulators of myogenesis and cannot proceed down the differentiation pathway. Instead, the NRMT1 knockout cells take on characteristics of C2C12 cells that have transdifferentiated into osteoblasts, including increased alkaline phosphatase (ALP) and type I collagen (Col1) expression and decreased proliferation rates ^30^. These data implicate NRMT1 as an important downstream target of CREB1 during myoblast differentiation, and suggest CREB1-mediated transcriptional regulation of NRMT1 as a conserved mechanism for governing the fate of many different stem cell lineages.

## Materials and methods

### Molecular cloning and mutagenesis

Fragments of the NRMT1 proximal promoter region were cloned into a modified luc2P/NF-κB-RE/Hygro vector, in which five copies of the NF-κB response element were removed, leaving only a minimal promoter. The 1,200 bp variants were generated by producing PCR products from HCT116 genomic DNA, and cloning into the luciferase vector using the Kpn1 and either HindIII or BglII restriction sites. Smaller truncations were generated by using the larger insert as a PCR template. Site-directed mutagenesis was utilized to generate point mutants in the putative TF binding sites (QuikChange system, Agilent Technologies, Santa Clara, CA). The following primers, and their reverse complements, were used to generate the point mutants: *SP1:* 5’-CGCGGGGGAGGGTTGAGCTGAGCGAGG-3’, *ETS:* 5’-GCCCGCCCCTAAACGCCCGTTAGTGACGTTGGCAGATCTG, *CRE/ATF:* 5’-CTAAACGCCCGGAAGTCGCGTTGGCAGATCTGG-3’. All constructs generated were verified by sequencing.

### Luciferase assays

HCT116 human colorectal carcinoma cells (1×10^3^) were plated on 96-well plates and allowed to adhere overnight. Cells were then co-transfected with 100 ng luciferase vector and 20 ng of pGL4-hRLuc-TK *(Renilla,* Promega, Madison, WI). 24 hours later, the Dual-Glo Luciferase System (Promega) was used to measure luciferase and renilla signals. The resulting signals were ratioed using a Cytation 5 Imaging Reader (BioTek, Winooski, VT). Empty luc2P/Hygro vector served as a negative control.

### DNA pulldowns

Nuclear lysates were produced from HCT116 cells as previously described ^31^. HCT116 cells were washed with ice cold PBS and lysed into PBS-I buffer (PBS, 0.5 mM PMSF, 25 mM β-glycerophosphate, 10 mM NaF). Cells were harvested by centrifugation, and following removal of supernatant the pellet was re-suspended in 2 package cell volume of buffer A (10 mM HEPES, pH 7.9, 1.5 mM MgCl2, 10 mM KCl, 300 mM sucrose, 0.5% NP40). Cells were incubated on ice for 10 minutes, harvested by centrifugation, and the cell pellet was re-suspended in 2/3 cell package volume of buffer B (20 mM HEPES, pH 7.9, 1.5 mM MgCl_2_, 420 mM NaCl, 0.2 mM EDTA, 2.5% glycerol). The mixture was sonicated for 5 seconds, and then centrifuged for 10 minutes at 10,400g. The supernatant was recovered and diluted isovolumetrically with buffer D (20mM HEPES, pH 7.9, 100 mM KCl, 0.2 mM EDTA, 8% glycerol). Buffers A, B, and D were all supplemented with an inhibitor cocktail (0.5 mM PMSF, 1 mM Na_3_VO_4_, 0.5 mM DTT, 1ug/mL leupeptin, 25 mM β-glycerophosphate, 10 mM NaF). Protein concentration of lysates was determined with the Pierce 660 Protein Assay Reagent (Thermo Scientific, Rockford, IL).

DNA pulldowns were performed as previously described with minor modifications ^32^. 100 bp oligos were synthesized (Invitrogen, Carlsbad, CA) with a biotin tag on the 5’ end of the forward strand. Oligos were annealed in 2x annealing buffer (20 mM Tris, pH 8.0, 100 mM NaCl, 2 mM EDTA), then used as baits in the HCT116 nuclear lysates. M-280 Streptavidin Dynabeads (Invitrogen) were used for pulldowns. Beads were washed once with PBS + 0.1% NP-40 and once with DNA binding buffer (DBB: 10 mM Tris, pH 8.0, 1 M NaCl, 0.05% NP40, 1 mM EDTA). 500 pmol of annealed oligos were diluted in 600 μL DBB and rotated for 30 minutes at 4°C. Following oligo binding, beads were washed once in DBB and twice with protein binding buffer (PBB: 50 mM Tris, pH 8.0, 150 mM NaCl, 0.25% NP40, 1 mM DTT, 1ug/mL leupeptin, 1ug/mL aprotinin, 0.5 mM PMSF). Nuclear extracts (300-500 μg) were added to beads in a final volume of 600 μL PBB, and rotated for 90 minutes at 4°C. Following incubation, beads were washed three times with PBB and twice with 1x PBS. Beads were re-suspended in 25 μl of Laemmli Sample Buffer (60 mM Tris pH 6.8, 2% SDS, 10% glycerol, 5% β-mercaptoethanol, 0.01% bromophenol blue). Samples were analyzed by Western blots as described below. Inputs represent 10 μg nuclear lysate. A list of oligo sequences used in this assay is included in Supplemental Table 1.

### Cell culture and small molecule treatment

HCT116 cells were cultured in McCoy’s 5A media (ThermoFisher, Grand Island, NY) supplemented with 10% fetal bovine serum (FBS; Atlanta Biologicals, Atlanta, GA) and 1% penicillin-streptomycin (P/S; ThermoFisher). C2C12 mouse muscle cells were cultured in Dulbecco’s Modified Eagle Medium (DMEM; ThermoFisher) with 10% FBS and 1% P/S. All cells were grown at 37°C and 5% CO_2_ on tissue culture treated plastic (Corning, Corning, NY). HCT116 and C2C12 cell lines were a generous gift from Dr. Ian Macara, Vanderbilt University.

For experiments involving CREB1 inhibition, HCT116 cells were treated with 10 nM of 666-15 (R&D Systems, Minneapolis, MN) for the indicated time. Control cells were treated with DMSO. For cell stress experiments, HCT116 cells were treated for 24 hours with 50 μM etoposide, 1 μM doxorubicin, 200 mM camptothecin, or 200 μM H_2_O_2_ (all Sigma, St. Louis, MO). Following treatment, cells were subsequently washed with phosphate buffered saline (PBS; Corning) and allowed to recover in fresh media for 6 hours before harvesting. For serum starvation experiments, HCT116 cells were washed with PBS, and then cultured in McCoy’s 5A media only for 24 hours. Following starvation, cells were released into full serum media for the indicated times. For experiments involving C2C12 muscle cell differentiation, cells were washed with PBS and then incubated in differentiation media (DMEM, 2% horse serum, 50 nM insulin (Sigma)) for the indicated times. Differentiation media was switched every 24 hours.

Cell viability experiments were carried out using the CellTiter 96 AQ_ueous_ One Solution Cell Proliferation Assay (Promega). For these experiments, 1×10^3^ cells were plated in triplicate in a 96-well plate. Daily cell number was determined by addition of 20 μL of AQ_ueous_ One Solution and measurement of absorbance at 490 nm. Fold change was calculated by dividing by absorbance measurements taken at day zero. Cells were re-fed at 48 hours.

### Western blots

Whole cell lysates were generated by lysing cells into buffer containing phosphatase inhibitors sodium fluoride (10 mM), β-glycerophosphate (1 mM) and sodium orthovanadate (1 mM, all Sigma). Protein samples were separated on 10 or 12% SDS-PAGE gels and transferred to nitrocellulose membranes (Bio-Rad, Hercules, CA) using a Trans-Blot SD Semi-Dry Transfer Cell (Bio-Rad). Membranes were incubated for one hour at room temperature in blocking buffer (5% w/v non-fat dry milk in TBS + 0.1% Tween 20 (TBS-T)). Primary and secondary antibodies were incubated in either 5% w/v non-fat dry milk or 3% w/v bovine serum albumin (BSA), Research Products International, Mount Prospect, IL) in TBS-T, depending on manufacturer’s instructions. Dilutions used for primary antibodies: mouse anti-Elk1 (E-5) (1:1000; sc-365876, Santa Cruz Antibodies, Dallas, TX), mouse anti-Elf1 (C-4) (1:1000; sc-133096, Santa Cruz), mouse anti-Ets-1 (C-4) (1:1000; sc-55581, Santa Cruz), mouse anti-GABP-α (G-1) (1:1000; sc-28312, Santa Cruz), mouse anti-CREB-1 (D-12) (1:1000; sc-377154, Santa Cruz), mouse anti-ATF1 (C41-5.1) (1:1000; sc-243, Santa Cruz), mouse anti-ATF2 (F2BR-1) (1:1000; sc-242, Santa Cruz), rabbit anti-SRF (D71A9) (1:1000; Cell Signaling Technology, Danvers, MA), rabbit anti-β-tubulin (9F3) (1:1000; Cell Signaling Technology), rabbit anti-GAPDH (1:3000; Trevigen, Gaithersburg, MD), rabbit anti-NRMT1 (1:1000; Chen, T et. al., 2007), rabbit anti-3meRCC1 (1:10,000; Chen, T et. al., 2007). Secondary antibodies used were donkey anti-mouse or donkey anti-rabbit (1:5000; Jackson ImmunoResearch, West Grove, PA). Blots were developed on a ChemiDoc Touch imaging system (Bio-Rad) using Clarity Western ECL Substrate (Bio-Rad) or SuperSignal West Femto Maximum Sensitivity Substrate (Thermo Scientific).

### Real time PCR analysis

HCT116 or C2C12 cells were lysed in TRIzol (Invitrogen) and mixed with chloroform to extract the RNA. RNA was pelleted in isopropanol, washed with ethanol, and resuspended in 100-150 μL sterile water. 1 μg RNA was synthesized into cDNA using the SuperScript First-Strand Synthesis System (Invitrogen). cDNA was diluted 1:10 in sterile water, and 2 μL of each sample was used for quantitative RT-PCR with 2x SYBR green Master Mix and the CFX96 Touch Real-Time PCR Detection System (Bio-Rad). Samples were run in triplicate. Transcript expression levels were determined using the ΔΔCT quantification method. Primer sequences (Invitrogen) were as follows: human NRMT1 5’-GCAGAGGTTTTTGAGGGAAG-3’ and reverse 5’-TTGGCTTGAACCAGGAAGTC-3’; human GAPDH forward 5’-ACAGCCTCAAGATCATCAGCAA-3’ and reverse 5’-CCATCACGCCACAGTTTCC-3’; human UBC forward 5’-CTGGAAGATGGTCGTACCCTG-3’ and reverse 5’-GGTCTTGCCAGTGAGTGTCT-3’; human c-Fos 5’-CTGGCGTTGTGAAGACCATG-3’ and reverse 5’-GGTTGCGGCATTTGGCTGC-3’; mouse GAPDH forward 5’-GGCAAATTCAACGGCACAGT-3’ and reverse 5’-CGCTCCTGGAAGATGGTGAT-3’; mouse HSP-90 forward 5’-AAAGGACTTGCGACTCGCC’3’ and reverse 5’-ATCAGCTCTGACGAACCCGA-3’; mouse NRMT1 forward 5’-TCTTCCCCCAGGTAGCTCT-3’ and reverse 5’-TGCAGAGGTTTTTAAGGGAAG-3’; mouse Pax7 forward 5’-CCGTGTTTCCCATGGTTGTG-3’ and reverse 5’-GAGCACTCGGCTAATCGAAC-3’; mouse myogenin forward 5’-CTACAGGCCTTGCTCAGCTC-3’ and reverse 5’-ACGATGGACGTAAGGGAGTG-3’; mouse MyoD forward 5’-GAGCACTACAGTGGCGACTC-3’ and reverse 5’-GCTCCACTATGCTGGACAGG-3’; mouse PPARγ forward 5’-GTGCCAGTTTCGATCCGTAGA-3’ and reverse 5’-GGCCAGCATCGTGTAGATGA-3’; mouse Sox9 forward 5’-GAGGCCACGGAACAGACTCA-3’ and reverse 5’-CAGCGCCTTGAAGATAGCATT-3’; mouse osteocalcin forward 5’-CCTGAGTCTGACAAAGCCTTCA-3’ and reverse 5’-GCCGGAGTCTGTTCACTACCTT-3’; mouse ALP 5’-TCAGGGCAATGAGGTCACATC-3’ and reverse 5’-CACAATGCCCACGGACTTC-3’; mouse type I collagen 1 5’-ATGCCTGGTGAACGTGGT-3’ and reverse 5’-AGGAGAGCCATCAGCACCT-3’; mouse Runx2 forward 5’-AAATGCCTCCGCTGTTATGAA-3’ and reverse 5’ - GCTCCGGCCCACAA-3’; and mouse osterix forward 5’-AGCGACCACTTGAGCAAACAT-3’ and reverse 5’-GCGGCTGATTGGCTTCTTCT-3’.

### Chromatin immunoprecipitation

HCT116 cells (7×10^6^) were plated one day before start of experiment. The following day, cells were either fixed (control) or serum-starved for 24 hours in McCoy’s 5A media. A subset of cells was then allowed to recover in complete media for 6 hours prior to fixation. Cells were cross-linked with 1% formaldehyde for 10 minutes at RT, and quenched with 1x glycine for 5 minutes at RT. The cell pellet was re-suspended in 600 μL ChIP lysis buffer (25 mM Tris-HCl, pH 7.5, 150 mM NaCl, 0.1% SDS, 5 mM EDTA, 1% Triton X-100, 0.5% sodium deoxycholate), and sonicated at 30% amplitude for 6 x 10 seconds. An input sample (10 ng/μL) was set aside for subsequent PCR amplification; 25 ng input was used in each reaction. Proteins were immunoprecipitated from precleared lysates overnight at 4°C using 4 μg of control IgG (rabbit or mouse, Cell Signaling Technology) or indicated antibody. Antibodies used were rabbit anti-phopsho-CREB1 Ser133 (87G3) (Cell Signaling Technology), mouse anti-Elk1 (E-5, sc-365876) (Santa Cruz Biotechnology), and rabbit anti-SRF (D71A9) XP (Cell Signaling Technology). Magnetic DynaBeads (Invitrogen) were pre-blocked in herring sperm DNA (Thermo Scientific) and subsequently added to lysates for 1 hour at 4°C. Depending on the antibody host species, either Protein A (rabbit) or Protein G (mouse) beads were used for pull-down. Following immunoprecipitation, beads were washed twice in low ionic strength ChIP dilution buffer (10 mM HEPES, pH 7.4, 50 mM NaCl, 10% glycerol, 1% NP-40), once in high salt ChIP wash buffer (500 mM NaCl, 20 mM Tris-HCl pH 8.0, 2 mM EDTA, 0.1% SDS, 1% NP-40), once in LiCl wash buffer (0.25 M LiCl, 10 mM Tris-HCl pH 8.0, 1 mM EDTA, 1% NaDoc, 1% NP-40) and twice in Tris-EDTA (TE). Two rounds of elution were performed with 100 μL elution buffer (1% SDS, 100 mM NaHCO3) for 15 minutes at RT. Samples were then de-cross-linked by adding 5 μL of 5 M NaCl and incubating overnight at 65°C. Samples were then treated for one hour at 45 °C with 1 μL of proteinase K (10 mg/ml stock; Millipore Sigma) and 1 μL of RNase A (24 mg/mL stock; Thermo Scientific). Eluted DNA was isolated using a Qiagen PCR purification kit (Qiagen, Hilden, Germany), and re-suspended in 30 μL of TE buffer. Equal amounts of DNA (2.5 μL) were amplified by PCR alongside input DNA and analyzed by running on 2% agarose gels. Primer sequences (Invitrogen) for the gene promoters examined were as follows: ICAM1 forward 5’-CGCCCGATTGCTTTAGCTTG-3’ and reverse 5’-GGCTGAGGTTGCAACTCTGA-3’; c-Fos forward 5’-GAGCAGTTCCCGTCAATCC-3’ and reverse 5’-GCATTTCGCAGTTCCTGTCT-3’; NRMT1 forward 5’-GTTAAATCCGGAGCCCGCG-3’ and reverse 5’-CGACCAGACGCGACAGCTC-3’; Egr1 forward 5’-TGCAGGATGGAGGTGCC-3’ and reverse 5’-AGTTCTGCGCGCTGGGATCTCTC-3’; CPLX1 forward 5’-GAACAGAGCAGCGCATATCA-3’ and reverse 5’-CTGGGTCAGACCTGAGCTTC-3’.

### Generation of C2C12 NRMT1 K/O line using CRISPR/Cas9

The CRISPR/Cas9 target site was the same location in the first coding exon as used for HCT116 cells ^33^, with the exception of a one nucleotide difference between the mouse and human genes. The following oligos were ordered from Invitrogen: top 5’-CACCGCCGAGCATGCCGTCCACCGT-3’, bottom 5’-AAACACGGTGGACGGCATGCTCGGC-3’. Cloning of oligos into the pSpCas9(BB)- 2A-Puro plasmid and generation of successfully transfected subclones was performed largely as before. Subclones were selected for further expansion following analysis for NRMT1 expression and N-terminal methylation by Western blot. A control cell line was generated in parallel by transfecting cells with an empty pSpCas9(BB)-2A-Puro plasmid and performing subsequent puromycin selection.

### Immunostaining

After 144 hours in differentiation media (DM), cells were fixed with 4% paraformaldehyde, permeabilized and immune-stained with antibodies against embryonic myosin heavy chain (F1.652, Developmental Studies Hybridoma Bank, Iowa City, Iowa). Cells were also co-stained with Hoechst (AnaSpec, Fremont, CA).

### Statistical analyses

All statistical analysis was performed using Prism 6 software (GraphPad, San Diego, CA). Statistical test used is denoted in figure legends. Results shown are mean ± standard deviation or standard error of the mean, as noted in figure legends.

## Results

### Identification of the NRMT1 transcription start site and minimal promoter region

To begin characterizing the transcriptional regulation of NRMT1, we first used the Ensembl and UCSC databases to help identify its transcription start site (TSS). The fulllength NRMT1 gene encompasses four exons, but only exons 2-4 are translated. There are 11 reported splice variants for NRMT1, but only four are predicted to encode the fulllength isoform (Ensembl database). Using the predicted Ensembl variant sequences, we cloned 1,200 base pairs (bp) of sequence immediately upstream of each of these four transcript variant’s predicted TSS into a luciferase vector containing a minimal promoter. We found that variant #2 produced a 61–fold increase in luciferase activity compared to empty vector control (Figure 1(a)). By comparison, variants #1 and #3 show little transcriptional activity over empty vector control, while variant #5 produced 35% of the activity of variant #2 (Figure 1(a)). These data indicate that variant #2 is the predominantly expressed form of NRMT1.

**Figure 1.**
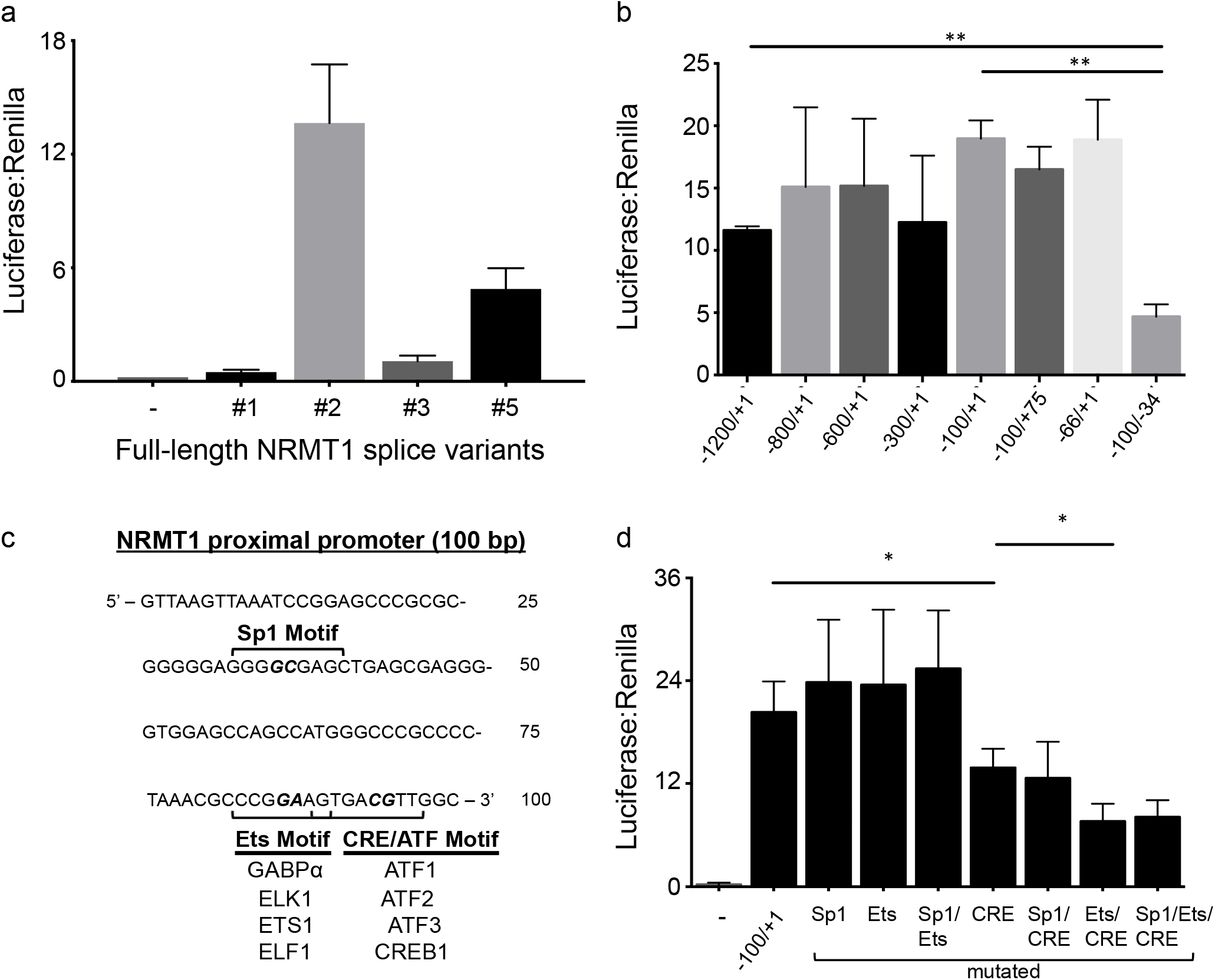
NRMT1 transcription is driven by its proximal promoter region. (a) The 1200 base pairs upstream of NRMT1 variant #2 produce more luciferase activity than the 1200 bp upstream of variants #1, 3, and 5, indicating variant #2 is the predominantly expressed form of NRMT1. (b) Compared to the full 1200 bp upstream of the variant #2 transcriptional start site (−1200/+1), removing 400 (−800/+1), 600 (−600/+1), 900 (−300/+1) or 1100 (100/+1) base pairs does not significantly decrease luciferase activity nor does removing the Sp1 binding motif (−66/+1). Addition of 75 bp after the transcription start site (−100/+75) also does not alter luciferase activity, but removal of the Ets and CRE/ATF motifs (−100/-34) causes a significant decrease. (c) Map of the 100 bp NRMT1 proximal promoter, including the Sp1, Ets, and CRE/ATF motifs. Listed are the family members shown to bind the sites in cells. (d) Single mutation of only the CRE site significantly decreases luciferase expression, while a double Ets/CRE mutation produces an additional decrease. A triple Sp1/Ets/CRE mutation produces no additional effect. * denotes p<0.05 and ** denotes p<0.01 as determined by unpaired t-test. n=3. Error bars represent ± standard deviation.

To better define the core promoter region of NRMT1, we next generated a series of serial truncation mutants in our luciferase vector. The 1,200 bp sequence immediately upstream of the NRMT1 TSS was used as a baseline and designated −1200/+1, with +1 representing the TSS. Removing up to 900 bp of upstream promoter sequence (−300/+1) did not significantly change transcriptional levels, and the highest amount of transcriptional activity actually was observed with the −100/+1 truncation, indicating that the proximal promoter region primarily drives NRMT1 transcription (Figure (1b)). Extending the promoter sequence 75 bp past the TSS (−100/+75) also did not change transcription levels (Figure (1b)). We conclude that the 100 bp proximal promoter region is the main regulatory unit promoting NRMT1 gene transcription.

### Identification of regulatory transcription factors

Analysis of the proximal promoter sequence using the PROMO program ^34^ revealed a number of predicted transcription factor (TF) binding sites in this region, including Sp1, the Ets family (28 members), and the CRE/ATF family (14 members) (Figure (1c)). Based on location of these TF binding sites, we generated two additional truncations within the proximal promoter region. One truncation (−66/+1) removed the Sp1 binding site, while the other truncation (−100/-34) removed the overlapping Ets and CRE/ATF sites. When these promoter truncations were tested in luciferase assays, only the truncation lacking the Ets and CRE/ATF sites caused a drop in gene transcription (Figure (1b)).

To determine whether the Ets or CRE/ATF binding site was primarily responsible for driving transcription, we next generated validated point mutants in the consensus binding sites for Ets *(GGAA* to G*TT*A) or CRE/ATF (T*GA*CGT to T*CG*CGT) ^35^ and tested them in luciferase assays (Figure 1(d)). As an additional control, we generated a point mutant in the Sp1 binding site (GGG*GC*GAGC to GGG*TT*GAGC) ^36^. Whereas point mutations in the Sp1 or Ets binding site did not decrease luciferase expression, disruption of the CRE binding site markedly reduced transcription (Figure 1(d)). Combining the CRE mutation with the Sp1 mutation did not further reduce luciferase signal (Figure 1(d)). However, disrupting both the CRE and Ets binding sites did lower transcription compared to disrupting CRE alone (Figure 1(d)). A triple mutation that included the Sp1 binding site did not further reduce transcription levels (Figure 1(d)). Therefore, we conclude that the overlapping Ets and CRE/ATF motifs drive transcription from the NRMT1 proximal promoter, with the CRE/ATF site predominating.

Combined, the Ets and CRE/ATF families contain more than forty unique transcription factors ^37,38^. Some of these factors have been shown to bind indiscriminately, where others require a specific sequence ^39^. To gain a better understanding of how the NRMT1 promoter is engaged, we began by utilizing the ENCODE ChIP-seq database to identify transcription factors from these families that are reported to bind within the NRMT1 proximal promoter region in cells. Within the CRE/ATF family, ENCODE reported that CREB1, ATF1, ATF2, and ATF3 showed strong binding to this region of DNA. Within the Ets family, GABPα, ETS1, ELK1, and ELF1 were reported to bind (denoted in Figure 1(c)).

Based on this information, we utilized a DNA pull-down assay to determine which of these TFs were capable of directly binding the NRMT1 proximal promoter sequence. 100 bp biotinylated oligos containing promoter sequence variations were incubated in HCT116 nuclear lysates. The oligos were retrieved using streptavidin-coated beads, and transcription factor binding was determined by Western blot. Among candidate CRE/ATF family members, only CREB1 was capable of binding the NRMT1 promoter sequence, but it was only capable of binding when the 100 bp DNA sequence was shifted 20 base pairs past the TSS (Figure 2(a)). This suggests that the TF dimer formed by CREB1 requires additional sequence past the TSS to serve as a platform for docking. ATF1, which belongs in the same TF subfamily as CREB1 and demonstrates some functional redundancy *in vivo* ^40,41^, had lower expression in the lysates and failed to bind any of the oligos (Figure 2(a)), demonstrating binding specificity at the NRMT1 promoter. ATF2, despite being detected in the nuclear lysates at levels comparable to CREB1, also failed to bind to any oligos over background levels (Figure 2(a)). We were unable to detect any ATF3 expression in the lysates (data not shown).

**Figure 2.**
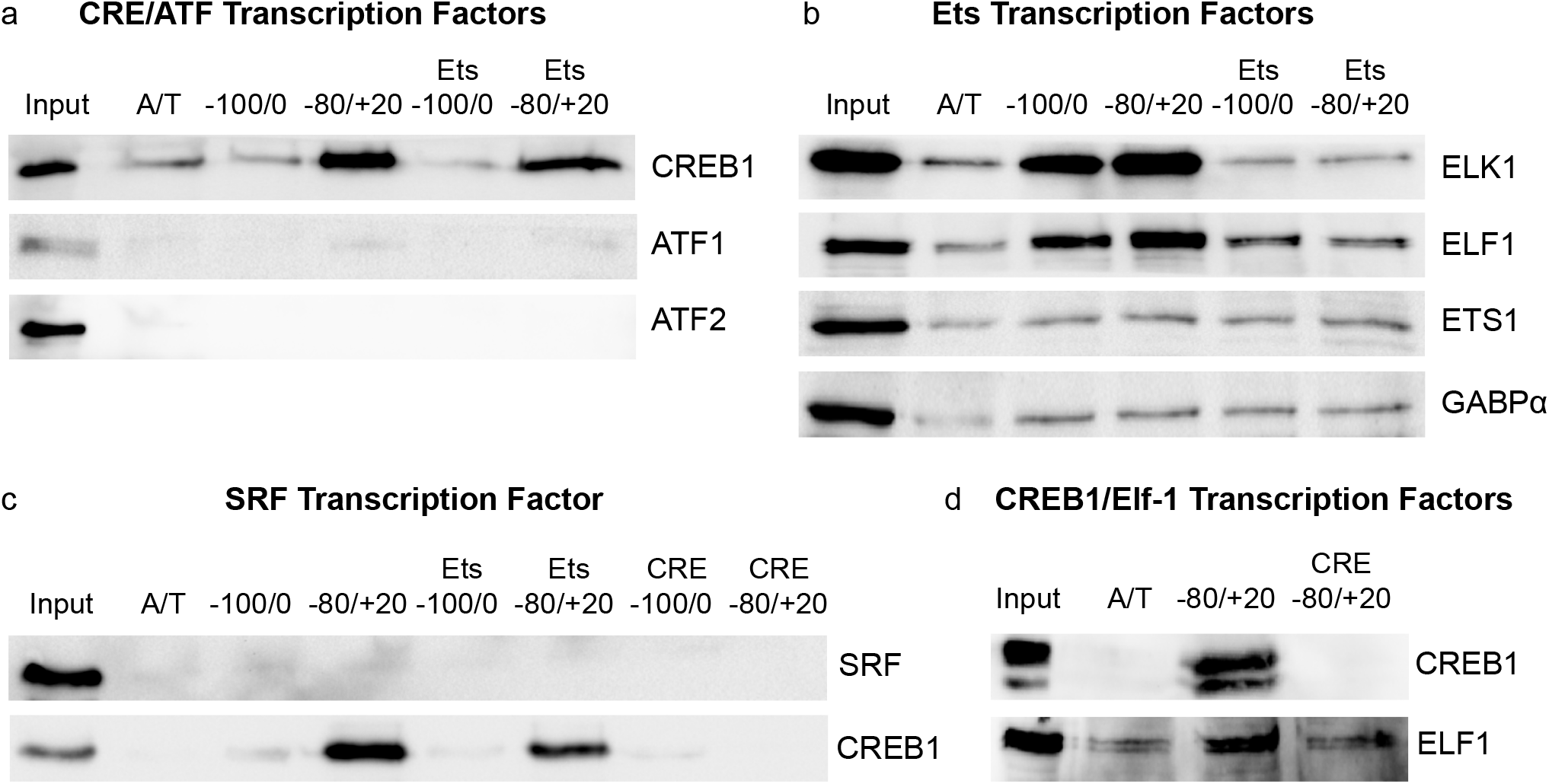
*In vitro* DNA binding assays show CREB1, ELK1, and ELF1 bind the NRMT1 proximal promoter. (a) Of the tested CREB/ATF family members, only CREB1 bound the NRMT1 proximal promoter sequence *in vitro*, though 20 bp past the transcriptional start site (−80/+20) was required for docking. Mutation of the Ets binding site (Ets −100/0, Ets −80/+20) did not affect CREB1 binding. (b) Of the Ets family members, only ELK1 and ELF1 bound the NRMT1 proximal promoter sequence above background levels. Addition of 20 bp past the transcriptional start site had no effect on ELK1/ELF1 binding, but mutation of the Ets binding site reduced binding to background levels. (c) Serum response factor, which often binds cooperatively with ELK1, does not bind the NRMT1 promoter. (d) Mutation of the CRE motif abolishes CREB1 binding to the NRMT1 promoter and also decreases EFL1 binding, indicating CREB1 recruits ELF1.

Ets family members also showed binding specificity, as neither ETS1 nor GABPα bound the 100 bp promoter sequence above background levels but both ELK1 and ELF1 showed robust binding (Figure 2(b)). In contrast to CREB1, shifting the promoter sequence 20 base pairs past the TSS did not significantly alter ELK1 or ELF1 binding, but binding was abolished by a point mutation in the core consensus Ets binding site (Figure 2(b)). ELK1 is a member of the TCF subfamily of the Ets family and is often functionally linked with a co-factor, serum response factor (SRF) ^42^. SRF binds to the serum response element (SRE), a palindromic DNA sequence that partially resembles the sequence immediately upstream of the ELK1 binding site within the NRMT1 promoter. Therefore, we also tested for SRF binding in our DNA pull-down assay. The NRMT1 promoter sequence was unable to bind SRF, even though SRF protein was detected in the nuclear lysate from cells cultured in growth media (Figure 2(c)).

Adjacent transcription factors can often demonstrate binding cooperativity, so we asked whether such binding existed at the NRMT1 proximal promoter. Mutating the Ets binding site, which eliminates binding of either ELK1 or ELF1, does not significantly reduce CREB1 binding (Figures 2(a) and 2(c)). Conversely, mutating the CRE/ATF site, which eliminates nearly all CREB1 binding, partially reduces the binding of ELF1 (Figure 2(d)). We conclude that CREB1, ELK1, and ELF1 are capable of biding the NRMT1 promoter in a non-competitive manner. However, as mutation of the Ets binding site does not significantly affect expression from the NRMT1 proximal promoter, we conclude that CREB1 is the main transcriptional driver of NRMT1 expression with some synergistic activation from ELK1/ELF1. Such synergistic actions between CREB1 and Ets family members have been previously reported at other target promoters ^43,44^.

### Conditional upregulation of NRMT1 expression

As CREB1 is an important transcriptional regulator of the DNA damage response, the cell cycle, and stem cell differentiation ^24,26,28,45,46^, we next wanted to determine what conditions stimulate CREB1-mediated activation of NRMT1 expression. Loss of NRMT1 sensitizes cells to both double-strand DNA breaks and oxidative damage ^2,3^, so we first treated cells with doxycycline, etoposide, camptothecin, or hydrogen peroxide and measured NRMT1 expression levels before and after treatment. No significant change in NRMT1 mRNA levels was seen with either DNA damage or oxidative stress (Figure 3(a)).

**Figure 3.**
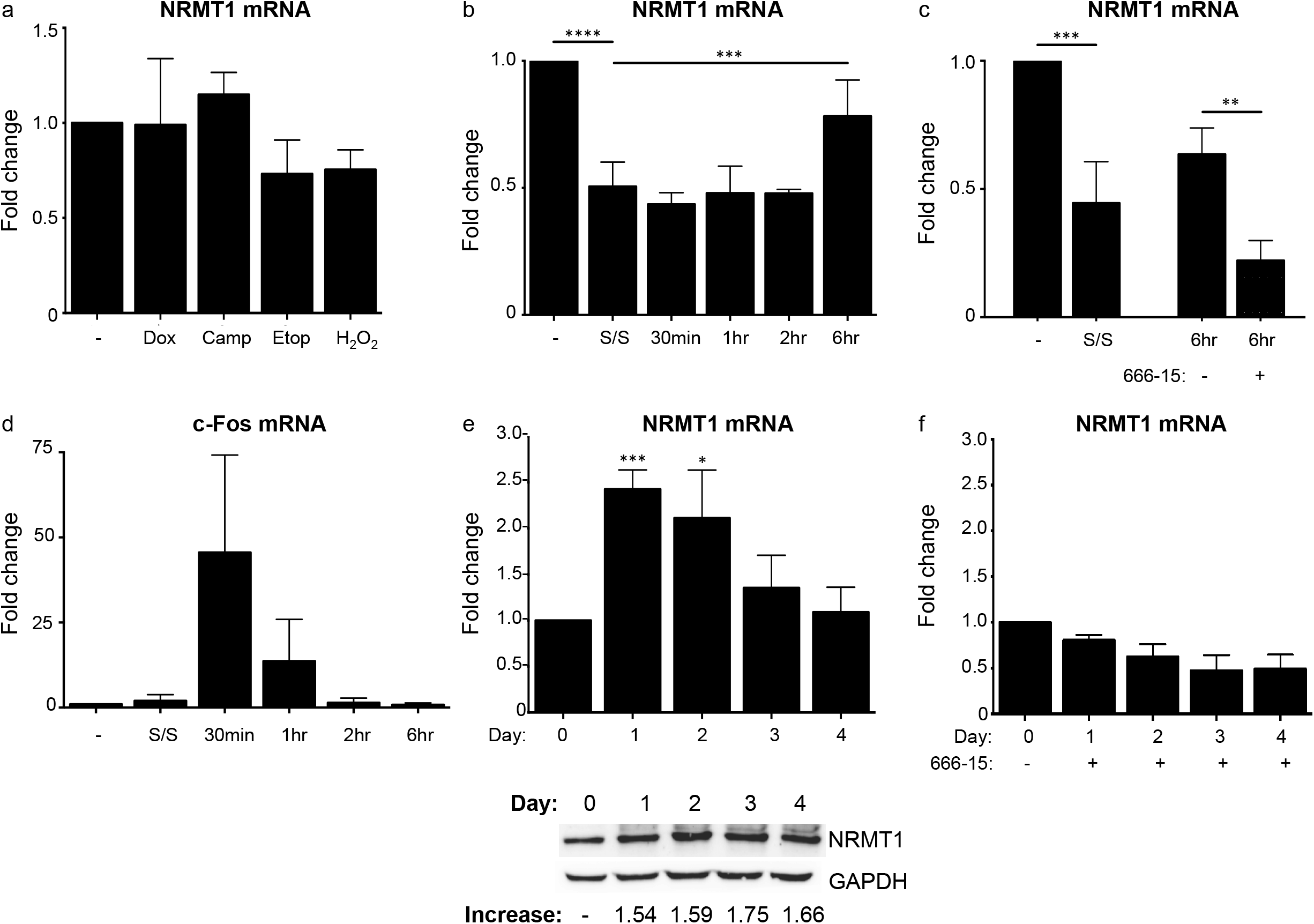
CREB1-mediated NRMT1 mRNA upregulation is stimulated during recovery from serum starvation and muscle cell differentiation. (a) Neither doxycycline (Dox), camptothecin (Camp), etoposide (Etop) nor hydrogen peroxide (H_2_O_2_) stimulate NRMT1 mRNA expression. (b) Removal from cell cycle by serum starvation (S/S) decreases NRMT1 mRNA levels but they are restored by 6 hr post-serum recovery. **** denotes p<0.0001, *** p<0.001 as determined by one-way ANOVA. n=4. (c) Treatment with the CREB1 inhibitor 666-15 prohibits restoration of NRMT1 mRNA levels 6 hr post-recovery. *** denotes p<0.001, ** p<0.01 as determined by unpaired t-test. n=4. (d) The delayed NRMT1 mRNA recovery after serum reintroduction is in contrast to expression levels of the immediate early gene c-Fos, which increase directly after serum addition, indicating NRMT1 is not an immediate early gene. (e and f) NRMT1 mRNA and protein levels also increase in differentiating C2C12 myoblasts, and this increase is also inhibited by 666-15. *** denotes p<0.001, * p<0.05 as determined by unpaired t-test. n=3. Error bars represent ± standard deviation.

Next, we wanted to monitor how cell cycle entry and withdrawal affected NRMT1 expression. To test this, we serum starved HCT116 cells for 24 hours and measured NRMT1 levels compared to cells grown in full serum media. Following serum starvation, NRMT1 transcription was significantly reduced by 50% (Figure 3(b)). We next measured the amount of time it took for NRMT1 transcription to recover following serum re-introduction. NRMT1 mRNA levels remained unchanged for the first two hours of recovery time, but by 6 hours, transcription had significantly increased to pre-withdrawal levels (Figure 3(b)). To ask if this increase in transcription following serum reintroduction was CREB1-dependent, we treated cells with the CREB1 small molecule inhibitor 666-15 during the 6-hour recovery period. 666-15 disrupts the interaction between the KID activation domain of CREB1 and the KIX binding domain of CBP and prevents CREB1-mediated transcriptional activation ^47^. Treatment of cells for 6 hours with 100 nM 666-15 blocked the recovery of NRMT1 transcription following serum reintroduction (Figure 3(c)). The NRMT1 expression pattern during serum starvation stands in contrast with the CREB1 target c-Fos, an immediate-early gene (IEG) whose mRNA levels rapidly rise and decline following serum stimulation (Figure 3(d)) ^48^. This suggests that NRMT1 transcription responds to sustained signaling, rather than the transient signaling that drives c-Fos expression.

Finally, we used the C2C12 muscle cell differentiation system to study the changes in NRMT1 levels during myogenesis. C2C12 myoblasts were differentiated into myotubes over a four-day time period by replacing growth media with low-serum differentiation media, and NRMT1 mRNA and protein levels were measured each day. After one day in differentiation media, NRMT1 mRNA levels increased nearly 2.5-fold compared to Day 0 myoblasts (Figure 3(e)). mRNA levels remained elevated but gradually declined during the differentiation time course (Figure 3(e)). NRMT1 protein levels also rose after one day in differentiation media, and remained elevated for the duration of the time course (Figure 3(e)). As with serum starvation, the increase in NRMT1 mRNA expression seen during muscle cell differentiation was abolished with 666-15 treatment, verifying it is also CREB1-dependent (Figure 3(f)).

### Conditional promoter binding in vivo

Once we determined conditions that stimulated CREB1-mediated NRMT1 expression, we wanted to confirm increased CREB1 and ELK1 occupancy of the NRMT1 promoter *in vivo* using chromatin immunoprecipitation (ChIP) assays. In normal growth conditions (full serum media), activated CREB1 (pSer133) was detected on both the control c-Fos promoter and the NRMT1 proximal promoter but was absent from the negative control ICAM-1 promoter (Figure 4(a)). Following serum starvation, activated pCREB1 was no longer detected on either the c-Fos or NRMT1 promoters (Figure 4(a)), in agreement with published data showing a loss of CREB1 DNA binding activity following serum withdrawal ^49^. However, following reintroduction of full serum media for 6 hours, binding to both promoters was restored at elevated levels (Figure 4(a)).

**Figure 4.**
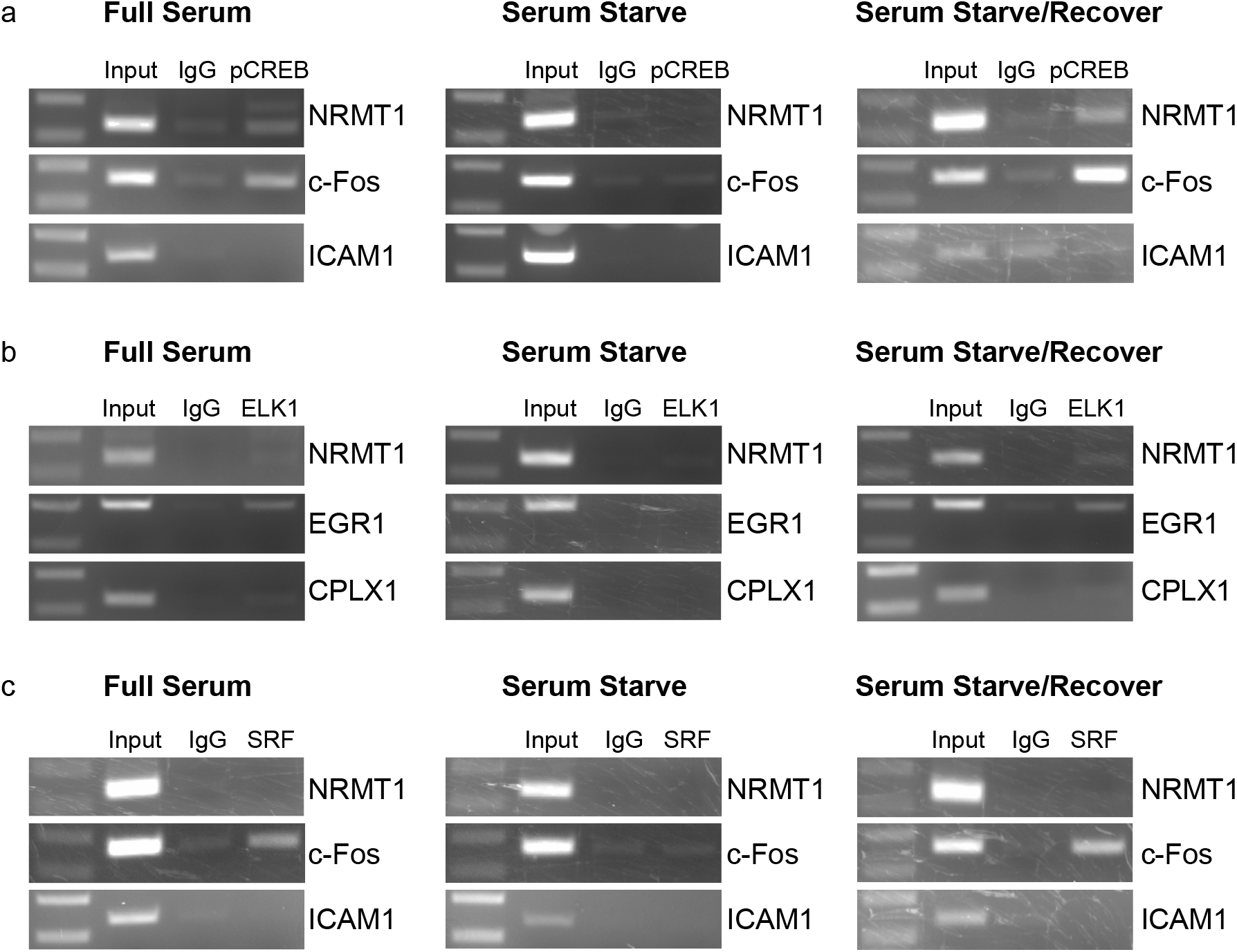
pCREB1 and ELK1 are recruited to the NRMT1 promoter in a serumdependent manner. (a) ChIP assay showing that activated CREB1 (pSer133) is bound to the NRMT1 promoter when HCT116 cells are cultured in growth media, and binding is lost during serum starvation. Activated CREB1 also binds its target c-Fos promoter during growth conditions, but not the negative control ICAM1. (b) ELK1 binds both the NRMT1 and target EGR1 promoters during periods of growth and recovery, but not during serum starvation. ELK1 fails to bind the negative control CPLX1. (c) SRF fails to bind the NRMT1 or ICAM1 promoters in any tested condition, despite binding its target c-Fos promoter in full serum media.

During normal growth conditions, ELK1 bound the control *Egr1* promoter as expected ^50^ and was also detected at a low level on the NRMT1 promoter (Figure 4(b)). Similar to pCREB1, ELK1 binding to the NRMT1 promoter was lost following serum starvation and restored following serum introduction (Figure 4(b)). In agreement with the DNA binding assay, SRF was unable to bind the NRMT1 promoter in any of the conditions tested, despite strongly binding the control c-Fos promoter in full serum media (Figure 4(c)). We conclude that pCREB1 and ELK1 are both capable of binding the NRMT1 promoter in cells, and this binding is regulated by cellular growth conditions.

### Effect of NRMT1 loss on muscle cell differentiation

Adult skeletal muscle represents a powerful system in which to study both quiescence and differentiation. Within skeletal muscle a small population of myogenic progenitor cells, named satellite cells, normally reside in a non-proliferative, quiescent state. When activated in response to exercise or injury, satellite cells exit quiescence, proliferate extensively as myoblasts, and differentiate and fuse to form myotubes and ultimately myofibers. To test if NRMT1 loss affects muscle cell differentiation, we first used CRISPR/Cas9 technology to deplete C2C12 cells of NRMT1 expression (Figure 5(a)). We differentiated the myoblasts into myotubes over a three-day time period by replacing growth media with low-serum differentiation media and used qRT-PCR to measure expression of markers of myoblast differentiation. The first striking observation was that even after 3 days in differentiation media, NRMT1 knockout (KO) C2C12 cells lacked expression of the master regulator of postnatal myogenesis, Pax7 (Figure 5(b)). The transcription factor Pax7 and its paralog Pax3 regulate myogenic precursor cell differentiation through their downstream targets, the myogenic regulatory factors, which include myogenic factor 5 (Myf5), myogenic differentiation 1 (MyoD), muscle-specific regulatory factor 4 (Myf4), and myogenin (MyoG) ^51^. Accordingly, MyoG expression was also minimally expressed over the differentiation period (Figure 5(c)), and NRMT1 KO C2C12 cells failed to fuse and produce myosin heavy chain (MHC) positive myotubes (Figure 5(d)).

**Figure 5.**
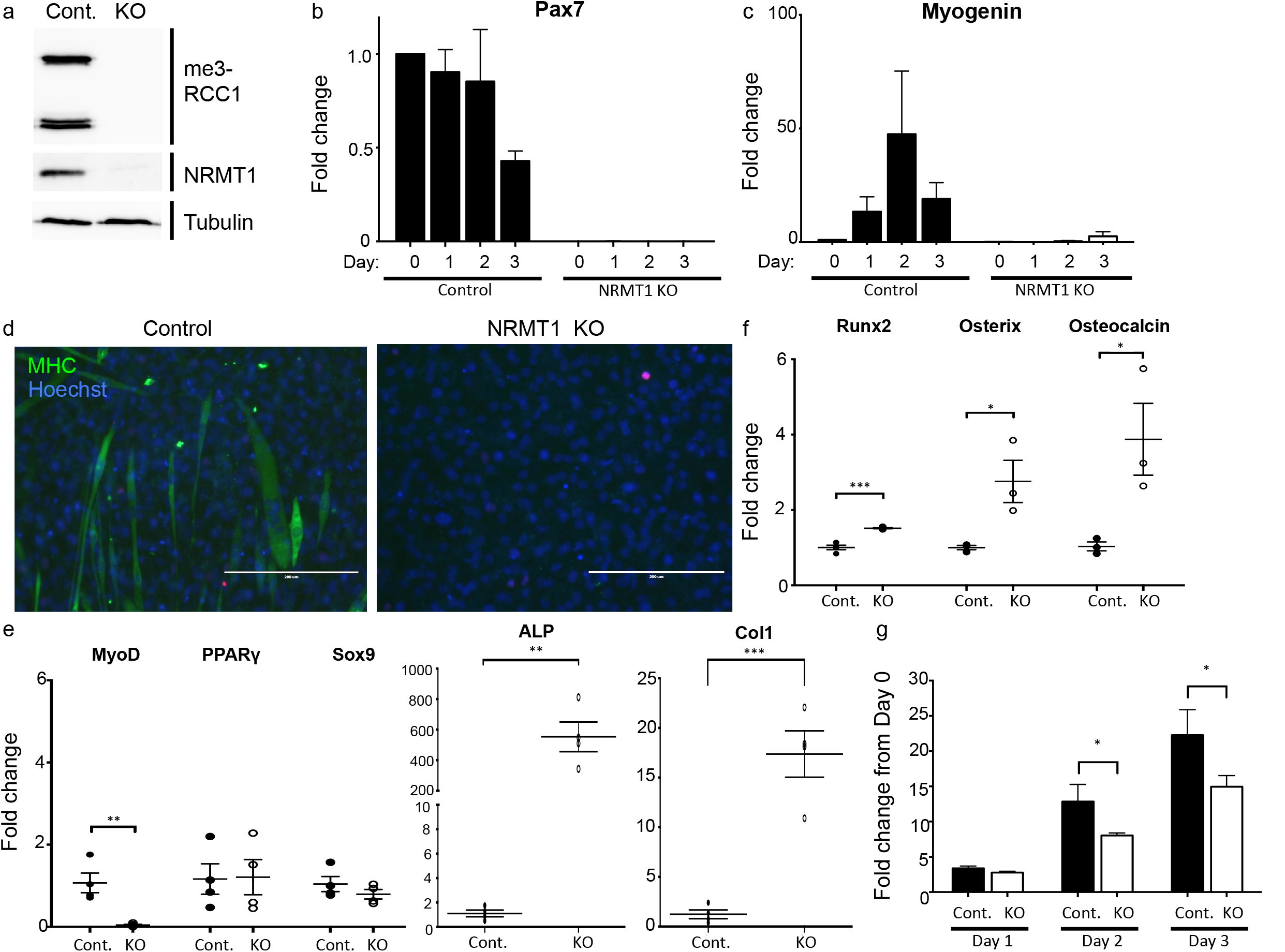
C2C12 cells lacking NRMT1 fail to properly differentiate and abnormally express osteogenic markers. (a) CRISPR/Cas9 technology was utilized to generate a NRMT1 knockout (KO) C2C12 cell line lacking NRMT1 expression and N-terminal trimethylation (me3-RCC1). Tubulin is used as a loading control. (b) Control C2C12 cells lose Pax7 expression as differentiation proceeds, but NRMT1 KO cells fail to express Pax7 mRNA at any time point. (c) The myogenin up-regulation seen in control cells during the differentiation time course is lost in KO cells. (d) After 6 days in differentiation media, NRMT1 KO cells fail to express myosin heavy chain (MHC) or form myotubes. Hoechst is used as a nuclear counterstain. Scale bars = 200 μm. (e) NRMT1 KO cells lose expression of the myogenic marker MyoD. However, while expression of adipogenic (*PPARƔ*) and chondrogenic (*Sox9*) markers remains unchanged, they have significantly increased expression of the osteogenic markers *alkaline phosphatase* (ALP) and *type 1 collagen* (Col1). (f) Expression of the osteogenic markers *Runx2, osterix,* and *osteocalcin* are also upregulated in NRMT1 KO cells. (g) NRMT1 KO cells grow slower than control C2C12 cells, another marker of osteogenic transdifferentiation. (e, f, g) *** denotes p<0.001, ** p<0.01, * p<0.05 as determined by unpaired t-test. n=3/4. Error bars represent ± standard error of the mean (e,f) or ± standard deviation (g).

As undifferentiated NRMT1 KO C2C12 cells lacked basal Pax7 expression, we wanted to determine if depletion of NRMT1 not only altered differentiation potential but cell type identity. Myoblasts are derived from a mesenchymal stem cell (MSC) progenitor pool that can also give rise to osteoblasts, adipocytes, and chondrocytes ^52^. To determine if loss of NRMT1 caused the C2C12 cells to transdifferentiate into bone, adipocyte, or chondrocyte lineages we performed qRT-PCR analysis on cells in normal growth media. As expected, there was little expression of MyoD further confirming the cells were no longer myogenic (Figure 5(e)). There was also little expression of PPARγ (a marker of the adipocyte lineage) or Sox9 (a marker of the chondrocyte lineage) (Figure 5(e)).

Interestingly, there was a significant increase in both ALP and Col1 expression in the NRMT1 KO cells, indicating they were transdifferentiating into an osteoblast lineage (Figure 5(e)). It has been previously shown that overexpression of osteogenic factors can drive C2C12 trans-differentiation into osteoblasts, increasing expression of osteoblastic markers and slowing C2C12 proliferation ^30^. To confirm similar transdifferentiation is happening with loss of NRMT1, we further assayed for *Runx2, osteocalcin* (OCN) and *osterix* (OSX) expression and measured the proliferation rates of the NRMT1 KO line. All three additional osteoblastic markers were increased in the NRMT1 KO line (Figure 5(f)), and their proliferation was significantly decreased from control C2C12 cells (Figure 5(g)). Taken together, these data demonstrate that NRMT1 is an important downstream target of CREB1 during myoblast differentiation and is a key component of the signaling pathways regulating mesenchymal stem cell lineage specification.

## Discussion

Though many important biological roles of NRMT1 have recently come to light, very little is known about its regulation. Its ubiquitous tissue expression and high levels of substrate methylation indicated a constitutive housekeeping gene ^1,8^. However, its subsequently identified roles in development and cell cycle regulation suggested a more intricate program of transcriptional regulation ^2,3,10^. Here we show for the first time that expression of NRMT1 is activated by the transcription factor CREB1 acting synergistically with members of the Ets family. We also demonstrate, as predicted, this transcriptional regulation of NRMT1 is closely tied to progression through the cell cycle.

CREB1 is a well-characterized transcriptional activator of genes that both induce and inhibit cell cycle progression, including cyclin A, c-Fos, RB, and Replication factor C3 (RFC3) ^45,48,53,54^. Given its expansive list of transcriptional targets, and the fact that many CREB1 targets are transcription factors themselves, the challenge is to now identify direct CREB1 targets and determine which ones are utilized in specific signaling pathways ^55^. This is especially important in MSCs, which give rise to a multipotent progenitor pool that can undergo osteogenesis, chondrogenesis, adipogenesis, or myogenesis ^52^. While activated CREB1 promotes the transcription of genes that drive muscle differentiation, including RB, Pax3, MyoD, and Myf5 ^53,56^, it also promotes the transcription of genes that drive osteogenesis, adipogenesis, and chondrogenesis ^57–59^. During osteogenesis, CREB1 activates genes that drive bone differentiation, including ALP and osteocalcin ^59,60^. During adipogenesis, CREB1 promotes the transcription of C/EBPβ and C/EBPδ, which in turn activate expression of PPARγ ^58^, and during chondrogenesis, CREB1 signaling drives Sox9 expression ^57,61^. How one transcription factor can promote such separate and distinct differentiation pathways of a common progenitor remains unknown.

Our data indicate that NRMT1 works downstream of CREB1 to specify differentiation pathways of the MSC multipotent progenitor pool. The presence of NRMT1 allows this pool to respond to myogenic signaling. However, in its absence, the myogenic transcriptional program is inhibited even in the presence of differentiation stimuli. N-terminal methylation by NRMT1 has been shown to promote protein/DNA interactions ^8^, and many known or predicted targets are themselves transcription factors ^1,4^. NRMT1 may fine-tune CREB1 signaling through activation of downstream, tissuespecific transcription factors. In this way, CREB1 would both increase transcription of its targets, and through NRMT1 expression, increase their activity as well.

We have recently shown that NRMT1 activates RB function in neural stem cells (NSCs) ^10^, and a similar pathway could be occurring in MSCs. While active RB is known to promote muscle differentiation ^62^, RB inactivation has been shown to promote osteoblast differentiation in culture ^63^. Similar to what is seen with loss of NRMT1, RB inactivation results in upregulation of *Runx2, OSX,* and *OCN* expression ^63^. This fits with a model of NRMT1 activating RB function in a MSC intermediate progenitor pool responding to myogenic signals. In the absence of NRMT1, RB becomes inactivated, promoting osteogenic differentiation under the same conditions. However, direct binding of RB to either the *Runx2* or *OSX* promoters could not be detected and both lack conventional E2F binding sites ^63^. Determining the direct targets of RB during myoblast and osteoclast differentiation will help test this model. The implicated role of RB downstream of NRMT1 in both MSCs and NSCs and the importance of CREB1 in both differentiation pathways ^28,53^, also posit that CREB1-mediated regulation of NRMT1 is conserved across stem cell types. Future experiments will also address this model.

Being able to therapeutically regulate MSC differentiation pathways would be a significant advance in regenerative medicine ^64^. The development of compounds that promote osteogenesis would be extremely beneficial in the treatment of osteoporosis ^60^. Conditions that promote osteoblastic programs over myoblastic programs, as seen with NRMT1 loss, have been shown to protect skeletal muscle from long-term denervation ^30,65^. Compounds that inhibit chondrocyte differentiation could slow the progression of osteoarthritis ^66^, and compounds that boost muscle stem cell differentiation would be invaluable for treatment of muscle degenerative diseases such as Duchenne Muscular Dystrophy ^67^. Both a better understanding of NRMT1 function and identification of other direct CREB1 targets responsible for fine-tuning CREB1 signaling during MSC differentiation will lead to novel ways of harnessing their potential as therapeutic agents.

## Acknowledgments

We thank Meghan Conner and Haley Parker for critical reading of the manuscript. This work was supported by a research grant form the National Institutes of Health to C.E.S.T [GM112721].

---

**Supplemental Table 1.**
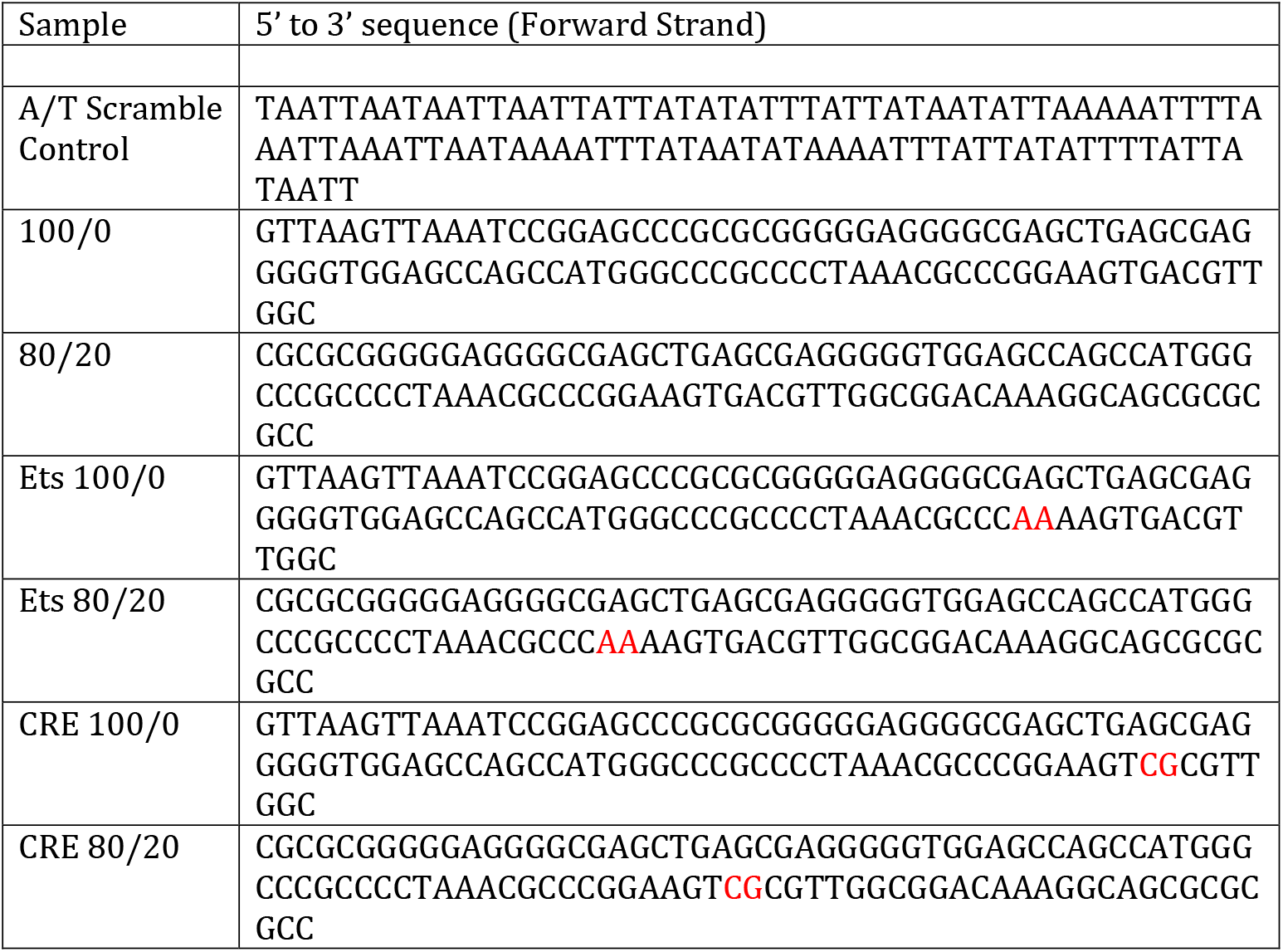
Oligos used in DNA pulldowns. Reverse strands are reverse complements of the forward strand oligo, but do not have a biotin tag attached. Mutations in Ets or CRE binding site are denoted in red.

## References

1 Tooley, C. E. et al. NRMT is an alpha-N-methyltransferase that methylates RCC1 and retinoblastoma protein. Nature 466, 1125–1128, doi:10.1038/nature09343 (2010).

2 Bonsignore, L. A., Butler, J. S., Klinge, C. M. & Schaner Tooley, C. E. Loss of the N-terminal methyltransferase NRMT1 increases sensitivity to DNA damage and promotes mammary oncogenesis. Oncotarget 6, 12248–12263. (2015).

3 Bonsignore, L. A. et al. NRMT1 knockout mice exhibit phenotypes associated with impaired DNA repair and premature aging. Mechanisms of Ageing and Development 146-148, 42–52, doi:http://dx.doi.org/10.1016/j.mad.2015.03.012 (2015).

4 Petkowski, J. J. et al. Substrate specificity of mammalian N-terminal alpha-amino methyltransferase NRMT. Biochemistry 51, 5942–5950, doi:10.1021/bi300278f (2012).

5 Bailey, A. O. et al. Posttranslational modification of CENP-A influences the conformation of centromeric chromatin. Proceedings of the National Academy of Sciences of the United States of America 110, 11827–11832, doi:10.1073/pnas.1300325110 (2013).

6 Cai, Q. et al. alpha-N-methylation of damaged DNA-binding protein 2 (DDB2) and its function in nucleotide excision repair. J Biol Chem 289, 16046–16056, doi:10.1074/jbc.M114.558510 (2014).

7 Dai, X. et al. Identification of novel alpha-n-methylation of CENP-B that regulates its binding to the centromeric DNA. Journal of proteome research 12, 4167–4175, doi:10.1021/pr400498y (2013).

8 Chen, T.et al. N-terminal alpha-methylation of RCC1 is necessary for stable chromatin association and normal mitosis. Nature cell biology 9, 596–603, doi:10.1038/ncb1572 (2007).

9 Nevitt, C., Tooley, J. G. & Schaner Tooley, C. E. N-terminal acetylation and methylation differentially affect the function of MYL9. The Biochemical journal 475, 3201–3219, doi:10.1042/bcj20180638 (2018).

10 Catlin, J. et al. Age-related neurodegeneration and cognitive impairments of NRMT1 knockout mice are preceded by misregulation of RB and expansion of the neural stem cell population. bioRxiv, 2021.2003.2015.435479, doi:10.1101/2021.03.15.435479 (2021).

11 Tammsalu, T. et al. Proteome-wide identification of SUMO2 modification sites. Science signaling 7, rs2, doi:10.1126/scisignal.2005146 (2014).

12 Larsen, S. C. et al. Proteome-wide analysis of arginine monomethylation reveals widespread occurrence in human cells. Science signaling 9, rs9, doi:10.1126/scisignal.aaf7329 (2016).

13 Wagner, S. A. et al. A proteome-wide, quantitative survey of in vivo ubiquitylation sites reveals widespread regulatory roles. Molecular & cellular proteomics: MCP 10, M111.013284, doi:10.1074/mcp.M111.013284 (2011).

14 Sharma, K. et al. Ultradeep human phosphoproteome reveals a distinct regulatory nature of Tyr and Ser/Thr-based signaling. Cell reports 8, 1583–1594, doi:10.1016/j.celrep.2014.07.036 (2014).

15 Olsen, J. V. et al. Quantitative phosphoproteomics reveals widespread full phosphorylation site occupancy during mitosis. Science signaling 3, ra3, doi:10.1126/scisignal.2000475 (2010).

16 Bade, D. et al. Modulation of N-terminal methyltransferase 1 by an N(6)-methyladenosine-based epitranscriptomic mechanism. Biochemical and biophysical research communications 546, 54–58, doi:10.1016/j.bbrc.2021.01.088 (2021).

17 Petkowski, Janusz J. et al. NRMT2 is an N-terminal monomethylase that primes for its homologue NRMT1. Biochemical Journal 456, 453–462. (2013).

18 Zhu, Y. et al. Discovery of coding regions in the human genome by integrated proteogenomics analysis workflow. Nat Commun 9, 903, doi:10.1038/s41467-018-03311-y (2018).

19 Wang, D. et al. A deep proteome and transcriptome abundance atlas of 29 healthy human tissues. Molecular systems biology 15, e8503, doi:10.15252/msb.20188503 (2019).

20 Bennett, M. L. et al. New tools for studying microglia in the mouse and human CNS. Proceedings of the National Academy of Sciences of the United States of America 113, E1738–1746, doi:10.1073/pnas.1525528113 (2016).

21 Clarke, L. E. et al. Normal aging induces A1-like astrocyte reactivity. Proceedings of the National Academy of Sciences of the United States of America 115, E1896–e1905, doi:10.1073/pnas.1800165115 (2018).

22 Alberini, C. M. Transcription factors in long-term memory and synaptic plasticity. Physiological reviews 89, 121–145, doi:10.1152/physrev.00017.2008 (2009).

23 Dyson, H. J. & Wright, P. E. Role of Intrinsic Protein Disorder in the Function and Interactions of the Transcriptional Coactivators CREB-binding Protein (CBP) and p300. The Journal of biological chemistry 291, 6714–6722, doi:10.1074/jbc.R115.692020 (2016).

24 Pregi, N., Belluscio, L. M., Berardino, B. G., Castillo, D. S. & Cánepa, E. T. Oxidative stress-induced CREB upregulation promotes DNA damage repair prior to neuronal cell death protection. Molecular and cellular biochemistry 425, 9–24, doi:10.1007/s11010-016-2858-z (2017).

25 Cataldi, A., di Giacomo, V., Rapino, M., Genovesi, D. & Rana, R. A. Cyclic nucleotide Response Element Binding protein (CREB) activation promotes survival signal in human K562 erythroleukemia cells exposed to ionising radiation/etoposide combined treatment. Journal of radiation research 47, 113–120, doi:10.1269/jrr.47.113 (2006).

26 Cheng, J. C. et al. CREB is a critical regulator of normal hematopoiesis and leukemogenesis. Blood 111, 1182–1192, doi:10.1182/blood-2007-04-083600 (2008).

27 Kim, J. M. et al. An activator of the cAMP/PKA/CREB pathway promotes osteogenesis from human mesenchymal stem cells. Journal of cellular physiology 228, 617–626, doi:10.1002/jcp.24171 (2013).

28 Mantamadiotis, T., Papalexis, N. & Dworkin, S. CREB signalling in neural stem/progenitor cells: recent developments and the implications for brain tumour biology. BioEssays: news and reviews in molecular, cellular and developmental biology 34, 293–300, doi:10.1002/bies.201100133 (2012).

29 Buckingham, M. & Relaix, F. PAX3 and PAX7 as upstream regulators of myogenesis. Semin Cell Dev Biol 44, 115–125, doi:10.1016/j.semcdb.2015.09.017 (2015).

30 Sondag, G. R. et al. Osteoactivin induces transdifferentiation of C2C12 myoblasts into osteoblasts. Journal of cellular physiology 229, 955–966, doi:10.1002/jcp.24512 (2014).

31 Wu, K. K. Analysis of protein-DNA binding by streptavidin-agarose pulldown. Methods Mol Biol 338, 281–290, doi:10.1385/1-59745-097-9:281 (2006).

32 Makowski, M. M. et al. An interaction proteomics survey of transcription factor binding at recurrent TERT promoter mutations. Proteomics 16, 417–426, doi:10.1002/pmic.201500327 (2016).

33 Shields, K. M. et al. Select human cancer mutants of NRMT1 alter its catalytic activity and decrease N-terminal trimethylation. Protein science: a publication of the Protein Society 26, 1639–1652, doi:10.1002/pro.3202 (2017).

34 Messeguer, X. et al. PROMO: detection of known transcription regulatory elements using species-tailored searches. Bioinformatics 18, 333–334, doi:10.1093/bioinformatics/18.2.333 (2002).

35 Pastorcic, M. & Das, H. K. An upstream element containing an ETS binding site is crucial for transcription of the human presenilin-1 gene. J Biol Chem 274, 24297–24307, doi:10.1074/jbc.274.34.24297 (1999).

36 Yang, G. et al. FoxO1 inhibits leptin regulation of pro-opiomelanocortin promoter activity by blocking STAT3 interaction with specificity protein 1. J Biol Chem 284, 3719–3727, doi:10.1074/jbc.M804965200 (2009).

37 Sharrocks, A. D. The ETS-domain transcription factor family. Nat Rev Mol Cell Biol 2, 827–837, doi:10.1038/35099076 (2001).

38 Hai, T. & Hartman, M. G. The molecular biology and nomenclature of the activating transcription factor/cAMP responsive element binding family of transcription factors: activating transcription factor proteins and homeostasis. Gene 273, 1–11, doi:10.1016/s0378-1119(01)00551-0 (2001).

39 Hai, T. W., Liu, F., Coukos, W. J. & Green, M. R. Transcription factor ATF cDNA clones: an extensive family of leucine zipper proteins able to selectively form DNA-binding heterodimers. Genes Dev 3, 2083–2090, doi:10.1101/gad.3.12b.2083 (1989).

40 Hummler, E. et al. Targeted mutation of the CREB gene: compensation within the CREB/ATF family of transcription factors. Proc Natl Acad Sci U S A 91, 5647–5651, doi:10.1073/pnas.91.12.5647 (1994).

41 Bleckmann, S. C. et al. Activating transcription factor 1 and CREB are important for cell survival during early mouse development. Mol Cell Biol 22, 1919–1925, doi:10.1128/mcb.22.6.1919-1925.2002 (2002).

42 Latinkić, B. V., Zeremski, M. & Lau, L. F. Elk-1 can recruit SRF to form a ternary complex upon the serum response element. Nucleic Acids Res 24, 1345–1351, doi:10.1093/nar/24.7.1345 (1996).

43 Yang, C., Shapiro, L. H., Rivera, M., Kumar, A. & Brindle, P. K. A role for CREB binding protein and p300 transcriptional coactivators in Ets-1 transactivation functions. Molecular and cellular biology 18, 2218–2229, doi:10.1128/mcb.18.4.2218 (1998).

44 Sawada, J. et al. Synergistic transcriptional activation by hGABP and select members of the activation transcription factor/cAMP response element-binding protein family. J Biol Chem 274, 35475–35482, doi:10.1074/jbc.274.50.35475 (1999).

45 Desdouets, C. et al. Cell cycle regulation of cyclin A gene expression by the cyclic AMP-responsive transcription factors CREB and CREM. Molecular and cellular biology 15, 3301–3309, doi:10.1128/mcb.15.6.3301 (1995).

46 Stewart, R., Flechner, L., Montminy, M. & Berdeaux, R. CREB is activated by muscle injury and promotes muscle regeneration. PLoS One 6, e24714, doi:10.1371/journal.pone.0024714 (2011).

47 Xie, F. et al. Identification of a Potent Inhibitor of CREB-Mediated Gene Transcription with Efficacious in Vivo Anticancer Activity. Journal of medicinal chemistry 58, 5075–5087, doi:10.1021/acs.jmedchem.5b00468 (2015).

48 Ramirez, S., Ait-Si-Ali, S., Robin, P., Trouche, D. & Harel-Bellan, A. The CREB-binding protein (CBP) cooperates with the serum response factor for transactivation of the c-fos serum response element. J Biol Chem 272, 31016–31021, doi:10.1074/jbc.272.49.31016 (1997).

49 Grimes, C. A. & Jope, R. S. CREB DNA binding activity is inhibited by glycogen synthase kinase-3 beta and facilitated by lithium. J Neurochem 78, 1219–1232, doi:10.1046/j.1471-4159.2001.00495.x (2001).

50 Vickers, E. R. et al. Ternary complex factor-serum response factor complex-regulated gene activity is required for cellular proliferation and inhibition of apoptotic cell death. Molecular and cellular biology 24, 10340–10351, doi:10.1128/mcb.24.23.10340-10351.2004 (2004).

51 Florkowska, A. et al. Pax7 as molecular switch regulating early and advanced stages of myogenic mouse ESC differentiation in teratomas. Stem Cell Res Ther 11, 238, doi:10.1186/s13287-020-01742-3 (2020).

52 Karantalis, V. & Hare, J. M. Use of mesenchymal stem cells for therapy of cardiac disease. Circulation research 116, 1413–1430, doi:10.1161/circresaha.116.303614 (2015).

53 Magenta, A. et al. MyoD stimulates RB promoter activity via the CREB/p300 nuclear transduction pathway. Molecular and cellular biology 23, 2893–2906, doi:10.1128/mcb.23.8.2893-2906.2003 (2003).

54 Chae, H. D., Mitton, B., Lacayo, N. J. & Sakamoto, K. M. Replication factor C3 is a CREB target gene that regulates cell cycle progression through the modulation of chromatin loading of PCNA. Leukemia 29, 1379–1389, doi:10.1038/leu.2014.350 (2015).

55 Impey, S. et al. Defining the CREB regulon: a genome-wide analysis of transcription factor regulatory regions. Cell 119, 1041–1054, doi:10.1016/j.cell.2004.10.032 (2004).

56 Chen, A. E., Ginty, D. D. & Fan, C. M. Protein kinase A signalling via CREB controls myogenesis induced by Wnt proteins. Nature 433, 317–322, doi:10.1038/nature03126 (2005).

57 Yokoyama, K. et al. Enhanced chondrogenesis of induced pluripotent stem cells from patients with neonatal-onset multisystem inflammatory disease occurs via the caspase 1-independent cAMP/protein kinase A/CREB pathway. Arthritis & rheumatology (Hoboken, N.J.) 67, 302–314, doi:10.1002/art.38912 (2015).

58 Lee, J. E., Cho, Y. W., Deng, C. X. & Ge, K. MLL3/MLL4-Associated PAGR1 Regulates Adipogenesis by Controlling Induction of C/EBPβ and C/EBPδ. Molecular and cellular biology 40, doi:10.1128/mcb.00209-20 (2020).

59 Huang, W. C. et al. Human osteocalcin and bone sialoprotein mediating osteomimicry of prostate cancer cells: role of cAMP-dependent protein kinase A signaling pathway. Cancer Res 65, 2303–2313, doi:10.1158/0008-5472.can-04-3448 (2005).

60 Zhang, Z. R. et al. Osthole Enhances Osteogenesis in Osteoblasts by Elevating Transcription Factor Osterix via cAMP/CREB Signaling In Vitro and In Vivo. Nutrients 9, doi:10.3390/nu9060588 (2017).

61 Zhang, Y., Huang, X. & Yuan, Y. Anti-inflammatory capacity of Apremilast in human chondrocytes is dependent on SOX-9. Inflammation research: official journal of the European Histamine Research Society ... [et al.] 69, 1123–1132, doi:10.1007/s00011-020-01392-4 (2020).

62 Huh, M. S., Parker, M. H., Scimè, A., Parks, R. & Rudnicki, M. A. Rb is required for progression through myogenic differentiation but not maintenance of terminal differentiation. The Journal of cell biology 166, 865–876, doi:10.1083/jcb.200403004 (2004).

63 Berman, S. D. et al. The retinoblastoma protein tumor suppressor is important for appropriate osteoblast differentiation and bone development. Molecular cancer research: MCR 6, 1440–1451, doi:10.1158/1541-7786.mcr-08-0176 (2008).

64 Ma, Z. J., Yang, J. J., Lu, Y. B., Liu, Z. Y. & Wang, X. X. Mesenchymal stem cell-derived exosomes: Toward cell-free therapeutic strategies in regenerative medicine. World journal of stem cells 12, 814–840, doi:10.4252/wjsc.v12.i8.814 (2020).

65 Furochi, H. et al. Overexpression of osteoactivin protects skeletal muscle from severe degeneration caused by long-term denervation in mice. The journal of medical investigation: JMI 54, 248–254, doi:10.2152/jmi.54.248 (2007).

66 van der Kraan, P. M. & van den Berg, W. B. Chondrocyte hypertrophy and osteoarthritis: role in initiation and progression of cartilage degeneration? Osteoarthritis and cartilage 20, 223–232, doi:10.1016/j.joca.2011.12.003 (2012).

67 Filippelli, R. L. & Chang, N. C. Empowering Muscle Stem Cells for the Treatment of Duchenne Muscular Dystrophy. Cells, tissues, organs, 1–14, doi:10.1159/000514305 (2021).

